# Beyond Exploration: Heightened Recent Reward Reliance and Midfrontal Sensitivity to Outcome Differences in ADHD

**DOI:** 10.64898/2026.11.09.765278

**Authors:** Clément Naveilhan, Remi Konan, Meggy Hayotte, Hervé Caci, Fabienne d’ Arripe-Longueville, Stephen Ramanoël

**Author notes:** CA, Corresponding Author: Stephen Ramanoel.

## Abstract

Adaptive decision-making requires balancing the exploitation of known rewards with the exploration of alternative options, yet how attention-deficit/hyperactivity disorder (ADHD) shapes this trade-off remains unclear. Using a restless four-armed bandit task and electroencephalography (EEG), we investigated the neural correlates of exploratory decisions and how recent rewards shape choice behavior and feedback processing in 23 adults with self- reported clinical ADHD diagnoses and 27 controls. Across 450 trials with dynamically changing payoffs, choices were classified against the highest last-observed payoff, and behavioral modeling quantified the influence of reward history on subsequent decisions. Contrary to our predictions, exploration rates, task performance, and neural differentiation between exploration and exploitation did not differ significantly between groups. However, adults with ADHD were more likely to repeat an option after high reward feedback and switch after low rewards, driven primarily by the immediately preceding outcome. At neural level, ADHD participants also showed greater midfrontal sensitivity to unsigned changes between consecutive payoffs, expressed in a feedback-evoked response spanning 280 to 568 ms and carried by slow activity below 2 Hz. Together, these findings dissociate the raw frequency of exploration from the dynamic weighting of recent outcomes in ADHD. Rather than reflecting an indiscriminate drive for novelty, behavior was characterized by heightened recent reward reliance paired with amplified slow-wave midfrontal sensitivity to outcome shifts. Crucially, this profile emerged without compromising overall performance, suggesting that, in this volatile environment, ADHD was associated with an increased weighting of recent outcomes rather than with a performance deficit.

## Introduction

Attention-deficit/hyperactivity disorder (ADHD) is a lifelong neurodevelopmental condition whose functional impact extends far beyond core symptoms of inattention, hyperactivity, and impulsivity (Faraone et al., 2024). In adulthood, ADHD is associated with reduced quality of life and substantial occupational, interpersonal, and financial difficulties (Brod et al., 2012; Joseph et al., 2019). To better explain these functional difficulties, it is necessary to understand how ADHD shapes decision-making processes, including evaluating past outcomes, weighing alternatives, and adapting to changing contingencies. Addressing these challenges requires identifying the neurocognitive mechanisms governing feedback integration and dynamic behavioral adjustment under uncertainty.

In this context, cognitive decision neuroscience leverages the exploration–exploitation dilemma, formalizing the fundamental tension between harvesting known rewards and gathering information about alternative options (Cohen et al., 2007). In non-stationary conditions such as restless multi-armed bandits where payoffs drift over time (Daw et al., 2006), successful adaptation requires balancing persistence on lucrative options against the discovery of improved alternatives. These explore–exploit trade-offs are modulated by frontal brain regions and dopaminergic tone, which contribute to calibrating value tracking versus exploratory sampling (Chakroun et al., 2020). Yet this exploratory sampling is not unitary: choices can emerge either through systematic, directed information-seeking or through random exploration (Dubois & Hauser, 2022; Wilson et al., 2014). In parallel, several findings from neurocomputational psychiatry propose that choice variability in ADHD stems from disrupted catecholaminergic tuning of neural gain, elevating stochastic decision noise during action selection (Hauser et al., 2016). Because random exploration is tightly linked to impulsivity (Dubois & Hauser, 2022), disentangling whether atypical choices in ADHD reflect altered value updating, intentional exploration, or elevated decision noise is essential to understanding the mechanisms underlying reward-guided behavior in this population.

Consistent with an overall bias away from exploitation, few emerging empirical evidence indicates that ADHD shifts the balance toward exploration. For example, adolescents with ADHD displayed greater exploratory switching during probabilistic reversal learning (Hauser, Iannaccone, Ball, et al., 2014). Similarly, adults with ADHD selected lower-value alternatives more frequently in a six-armed bandit task, and exploration was positively associated with hyperactive symptoms (Addicott et al., 2021). Yet the adaptive value of this exploratory drive depends on contextual reward dynamics. For example, in a foraging task with decaying yields, individuals with ADHD outperformed peers by abandoning depleted patches earlier (Barack et al., 2024). Heightened exploration can thus prove beneficial under resource depletion, but costly when the context of the task favors the sustained exploitation of stable options.

Beyond this explanation of contextual demands, altered reinforcement processing offers a complementary account for how ADHD biases this explore–exploit balance. Theoretical findings emphasize differences in the effects of reinforcement and the integration of rewards over time, rather than a uniform increase in reward sensitivity (Luman et al., 2010). The dopamine transfer deficit hypothesis proposes that behavior in ADHD depends more strongly on immediately delivered reinforcement because predictive cues acquire weaker reinforcing properties (Tripp & Wickens, 2008). Reward magnitude, frequency, and immediacy must nevertheless be distinguished, because sensitivity to one does not establish sensitivity to the others (Luman et al., 2009). Together these accounts motivate testing whether a recently obtained high reward more strongly promotes repetition of the rewarded choice in ADHD. Such an effect could coexist with an overall tendency toward exploration, and individuals might sample alternatives frequently but remain with an option after receiving a particularly rewarding outcome.

To maintain an optimal balance between feedback-driven persistence and exploratory switching, the brain recruits neural circuits that track outcome value and alternative opportunities. Alongside frontoparietal networks (Daw et al., 2006), the dorsal anterior cingulate cortex (dACC) activates more strongly during information gathering than reward exploitation (Blanchard & Gershman, 2018). In a simultaneous EEG–fMRI paradigm, trial-by- trial feedback-related negativity was linked to dACC activity, with its amplitude tracking surprise rather than reward prediction errors, to signal unexpected outcomes that warrant behavioral adjustment (Hauser, Iannaccone, Stämpfli, et al., 2014). Extending this circuit, Frank et al. (2015) showed that medial frontal–subthalamic mechanisms regulate decision caution under value conflict, where midfrontal theta dynamics interact with the subthalamic nucleus to elevate decision thresholds when competing options have similar learned values. Together, these studies motivate examining midfrontal electrophysiological responses during decision- making and feedback evaluation.

While fMRI maps the cerebral bases governing adaptive choice, electrophysiology is better suited to isolate the rapid processing that precedes and follows exploratory decisions. In particular, frontal midline activity dynamically indexes uncertainty and surprise across both decision modes in healthy adults (Cavanagh et al., 2012). Exploratory actions are accompanied by heightened choice-locked N200 and feedback-locked P300 components, reflecting response conflict during selection and deeper feedback re-evaluation (Hassall et al., 2019; Hassall & Krigolson, 2020), while trial-level reinforcement-learning signals predict subsequent choices (Cohen & Ranganath, 2007). Crucially, event-related potential (ERP) evidence dissociates reward magnitude from valence and expectancy, tracking both signed and unsigned prediction errors (Sambrook & Goslin, 2015; Yeung & Sanfey, 2004). Although such conflict- and feedback-related potentials are frequently altered in ADHD (Kaiser et al., 2020), electrophysiological studies of the exploration–exploitation trade-off have focused predominantly on neurotypical populations. Consequently, it remains unknown whether exploratory tendencies in adults with ADHD reflect altered deliberation during choice selection or differences in the downstream evaluation and integration of feedback.

To address this gap, we combined a four-armed restless bandit task with high-density EEG to examine exploration, feedback processing, and reward-recency effects in adults with ADHD and controls participants. We tested two primary hypotheses. First, we predicted that adults with ADHD would make more exploratory choices and exhibit greater ERP differentiation between exploration and exploitation over midfrontal electrodes, reflecting heightened deliberation or response conflict during action selection independent of raw choice frequency. Second, we predicted that adults with ADHD would be more strongly influenced by a recent reward, exhibiting an increased probability of repeating highly rewarded choices in association with an amplified midfrontal ERP contrast between high- and low-reward outcomes.

## Materials and Methods

### Participants

Fifty-four adults took part in the study, 30 control participants and 24 participants with a self- reported clinical diagnosis of ADHD. All participants were right-handed, between 18 and 40 years old, with normal or corrected to normal vision. All the participants were recruited from the local university and the surroundings and were compensated 20 euros for their participation. They all signed an informed consent form, and the study was approved by the local ethics committee (CER-2024-089). Based on the French version of the ASRS-v1.1 questionnaire (Caci et al., 2014; Kessler et al., 2005; Morin et al., 2016), which all participants completed prior to the experiment, three control participants were removed as they had exceeded the screening threshold of 4/6 in the Part A of the ASRS-v1.1 (respectively: 6/6, 5/6 and 4/6).

Each participant completed 450 trials of the four-armed restless bandit task in 3 blocks of 150 trials (**Figure 1.a**), with the payoffs of the four options drifting independently across trials according to a decaying Gaussian random walk (**Figure 1.b** and see Experimental design section), with a break between each block. To verify that participants engaged with the task rather than responding at random, we tested each participant against a participant-specific chance level. Because the payoffs available differed between participants, we simulated 10,000 agents that chose uniformly at random among the four options for the exact sequence of payoffs presented to that participant, and then expressed the resulting earnings as points per trial. One ADHD participant was excluded, having earned no more than this null distribution: 2,258 of the 10,000 random agents matched or exceeded this participant’s earnings (51.78 vs 51.40 points per trial, one-sided *p* = .226, **Figure 1.c**). All remaining participants earned more than their own null distribution (*z* = +10.4 to +24.4, all *p* < .0001).

**Figure 1.**
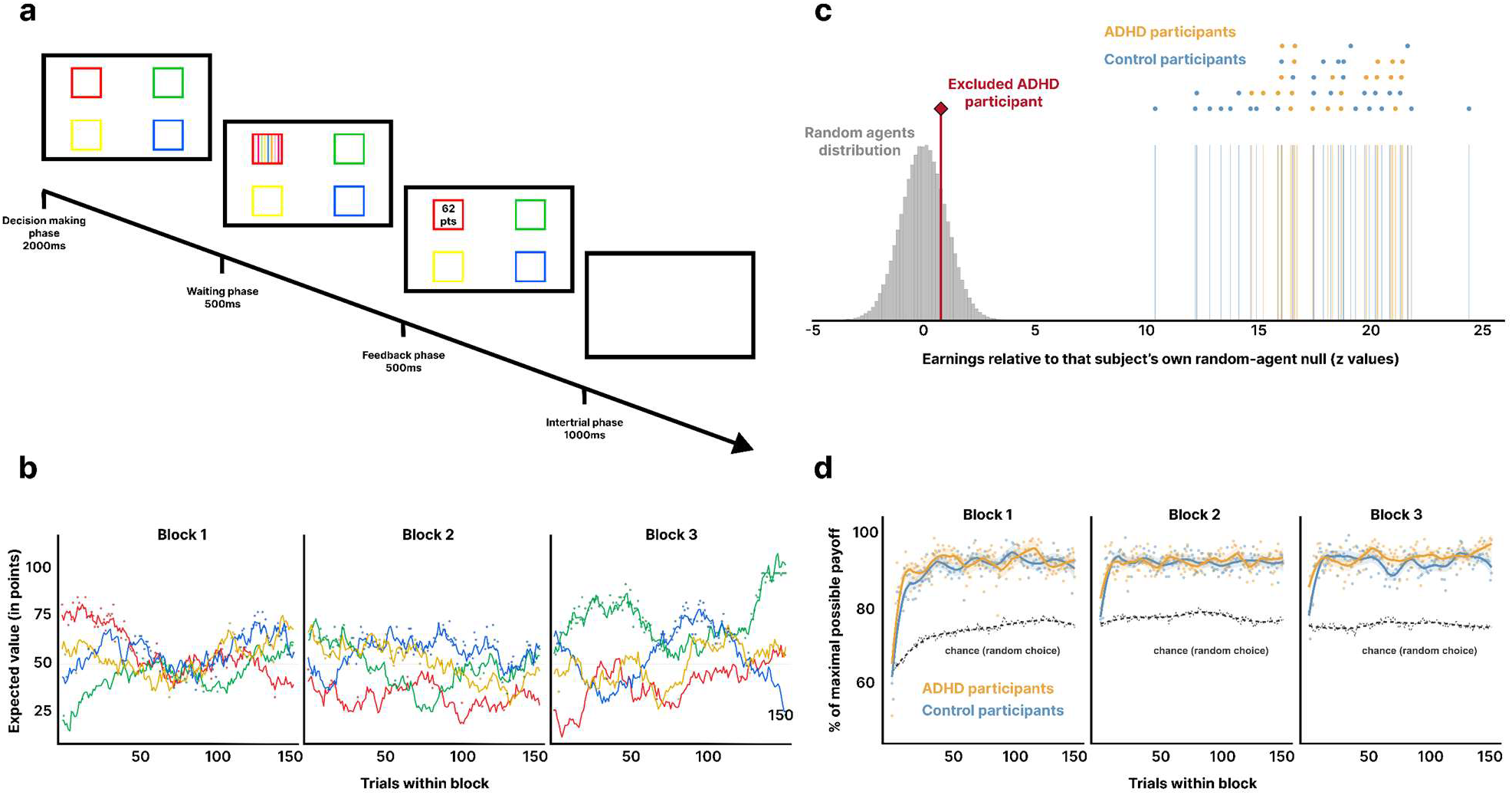
Presentation of the protocol. **(a)** Trial structure of the restless four-armed bandit. Participants had 2,000 ms to choose one of four options, followed by a 500 ms highlight of the chosen option, a 500 ms presentation of the payoff, and a 1,000 ms inter-trial interval. Three blocks of 150 trials were completed. **(b)** Reward environment for one representative participant (each participant’s specific values were independent). Colored lines show each option’s latent expected value, drifting as an independent mean-reverting Gaussian random walk, so no option remains optimal; these values were never shown. Dots show the payoff obtained on the chosen option (latent value + observation noise, SD = 4.0), tracing the choice sequence. **(c)** Performance-based exclusion. For each participant, earnings are expressed as a z score relative to 10,000 agents choosing randomly on that participant’s own payoff matrix (grey = pooled null). All retained participants scored z = +10.4 to +24.4 (*p* < .0001); one ADHD participant (red) scored z = +0.76 (*p* = .226, within chance) and was excluded. **(d)** Within-block learning. Points show the mean payoff obtained per trial relative to the best available payoff; lines are LOESS fits with 95% CIs. The dashed line marks chance (close to 75%, since a random chooser earns the four-option average while the denominator is the best option), which rises within a block as the walks converge; both groups start near or below chance and exceed it within a few trials.

After these exclusions the final sample size was 27 control participants (*M*_age_ = 23.7 SD = 3.0; 11 women) and 23 ADHD (*M*_age_ = 25.4 SD = 4.3; 12 women, 6.4 ± 6.9 years post clinical diagnosis of ADHD by clinician, (see **Table 1**). Control participants reported taking no psychotropic medication. Participants with ADHD who were taking medication were instructed to refrain from taking it for at least 48 h before the experimental session, a delay exceeding ten elimination half-lives of methylphenidate (Kimko et al., 1999), in order to avoid acute medication effects on decision-making (Agay et al., 2010). However, medication status, the specific compounds prescribed and adherence to this instruction were not systematically recorded.

**Table 1.**
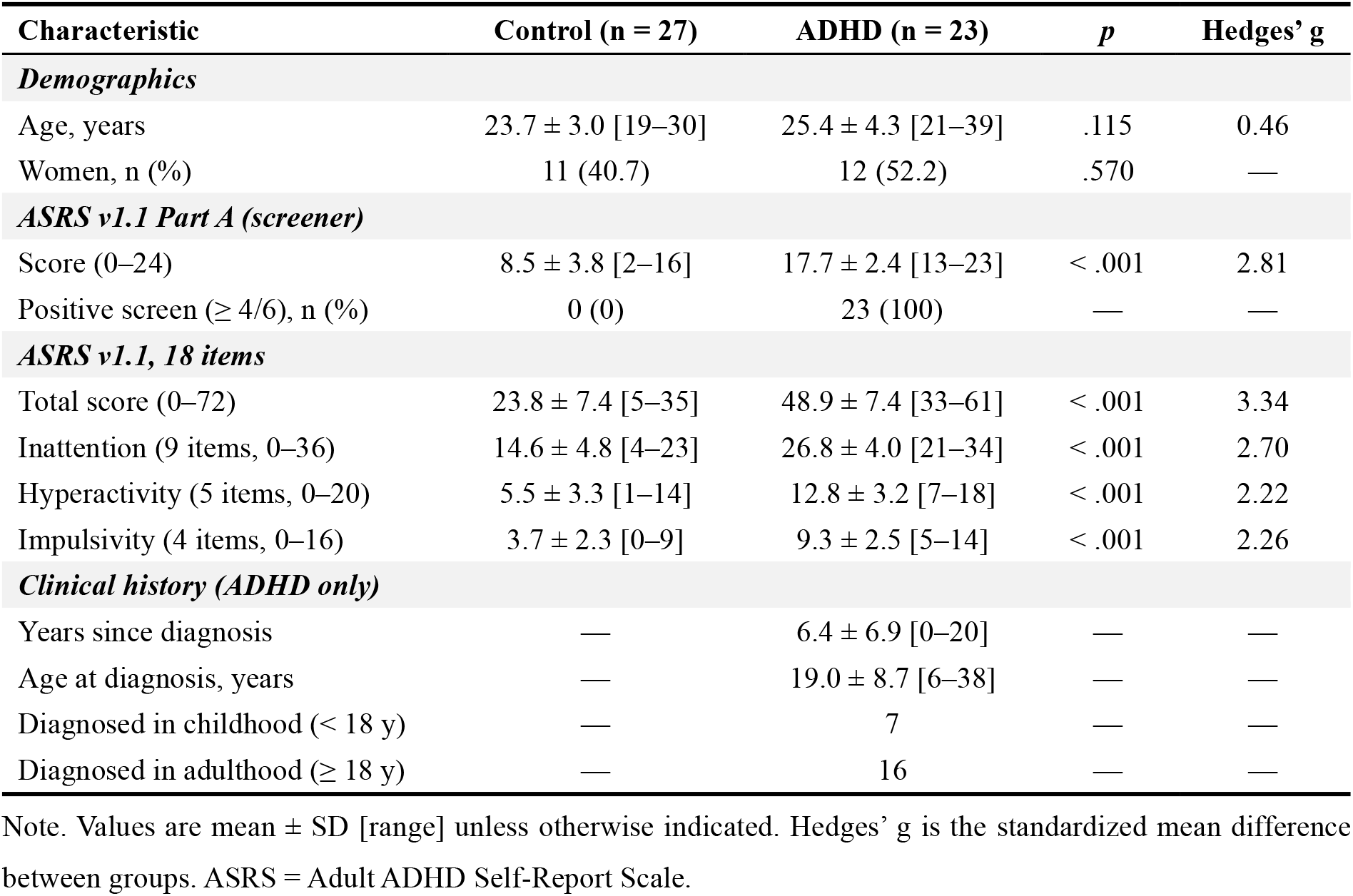
Participant characteristics (N = 50)

| **Characteristic** | **Control (n = 27)** | **ADHD (n = 23)** | ***p*** | **Hedges’ g** |
|---|---|---|---|---|
| ***Demographics*** |  |  |  |  |
| Age, years | 23.7 ± 3.0 [19–30] | 25.4 ± 4.3 [21–39] | .115 | 0.46 |
| Women, n (%) | 11 (40.7) | 12 (52.2) | .570 | — |
| ***ASRS v1.1 Part A (screener)*** |  |  |  |  |
| Score (0–24) | 8.5 ± 3.8 [2–16] | 17.7 ± 2.4 [13–23] | < .001 | 2.81 |
| Positive screen (≥ 4/6), n (%) | 0 (0) | 23 (100) | — | — |
| ***ASRS v1.1, 18 items*** |  |  |  |  |
| Total score (0–72) | 23.8 ± 7.4 [5–35] | 48.9 ± 7.4 [33–61] | < .001 | 3.34 |
| Inattention (9 items, 0–36) | 14.6 ± 4.8 [4–23] | 26.8 ± 4.0 [21–34] | < .001 | 2.70 |
| Hyperactivity (5 items, 0–20) | 5.5 ± 3.3 [1–14] | 12.8 ± 3.2 [7–18] | < .001 | 2.22 |
| Impulsivity (4 items, 0–16) | 3.7 ± 2.3 [0–9] | 9.3 ± 2.5 [5–14] | < .001 | 2.26 |
| ***Clinical history (ADHD only)*** |  |  |  |  |
| Years since diagnosis | — | 6.4 ± 6.9 [0–20] | — | — |
| Age at diagnosis, years | — | 19.0 ± 8.7 [6–38] | — | — |
| Diagnosed in childhood (< 18 y) | — | 7 | — | — |
| Diagnosed in adulthood (≥ 18 y) | — | 16 | — | — |

Note. Values are mean ± SD [range] unless otherwise indicated. Hedges’ g is the standardized mean difference between groups. ASRS = Adult ADHD Self-Report Scale.

An a priori power calculation was conducted for the directional hypothesis that exploration rates would be more elevated in ADHD. To this end we used the results from Hauser *et al*. (2014) who reported reduced decision steepness in ADHD (r = .360, approximate d = .77), although this model parameter differs slightly from the empirical exploration rates measured here. A one-sided independent t-test (α = .05) indicated that n = 22 per group (N = 44) achieved 81% power for d = .77. Despite this directional framing at the design stage, all the comparisons in the following were evaluated two-sided to ensure more conservative inference. With our final analyzed cohort (27 controls, 23 ADHD), the design provided approximately 76% power for d = .77 (80% power for d ≥ .81) under equal-variance assumptions. This sensitivity analysis demonstrates adequate power for large effects in the primary exploration-rate comparison, but does not extend to subtler effects or secondary measures. This limitation is acknowledged in the discussion.

### Experimental design

The decision-making was assessed with a four-armed restless bandit task (Daw et al., 2006), implemented from the Millisecond Software Inquisit task library and adapted for this study. The task was presented with Inquisit 7 on a 27-inch ASUS ROG Swift OLED monitor, ensuring millisecond accuracy between image presentation and EEG recordings (Dimigen & Stein, 2026). Responses were made on a Razer Huntsman Elite keyboard using the numeric keypad keys, with the right index finger only. The keys were individually illuminated in the colors of the corresponding on-screen boxes. To standardize the motor component of the decision, participants were required to return their finger to the center of the four keys before each new trial. The task was synchronized with the EEG recording via Lab Streaming Layer (Kothe et al., 2025), with markers sent at the onset of both the choice display and the payoff display. These two events were then used for the analyses centered on the decision making and the feedback processing phases, respectively.

This paradigm operationalizes the exploration–exploitation dilemma and allows us to observe directly the strategy by which participants optimize their choices. In each trial four colored boxes were displayed, each of which could return an integer payoff between 1 and 100 points, but only the payoff of the selected box was revealed, so feedback was partial and information about the other three options could only be obtained by sampling them (**Figure 1.a**). The payoffs of the four options drifted independently across trials, following a decaying Gaussian random walk (**Figure 1.b)**: in each trial the latent mean of an option was adjusted slightly towards a central value of about 51 points and then perturbed by Gaussian noise, and the payoff shown to the participant was drawn around that latent mean with a further, well-behaved Gaussian noise. The drift and observation parameters recovered from the logged payoff sequences (diffusion standard deviation 2.79 points, observation standard deviation 4.02 points, decay towards the center close to unity) matched those of the original Daw et al. (2006) implementation (2.8 and 4). Because the payoffs keep drifting, no single option remains optimal indefinitely, and continued exploration is required throughout the task. Participants had 2,000 ms to select a box, and if no response was made within that window a red cross was displayed to indicate a missed trial and the next trial began. Missed trials were rare (27 trials in total across all participants). Following the 2000ms of fixed response time, a fixed waiting screen appeared displaying colored bars for 500 ms, after which the payoff obtained was displayed for 500 ms, followed by a 1,000 ms blank screen separating consecutive trials, leading to trials lasting 4 s. These timing parameters were chosen to accommodate the attentional difficulties of participants with ADHD while maximizing the number of trials available for the EEG analyses. **Figure 1.d** shows that participants started below chance on the first trial of Block 1 (dashed line around 75%, since a random chooser earns the four-option average relative to the best option), rose above it by around trial 5, and then remained stable. In Blocks 2 and 3, performance began each block already above chance, since the reward environment kept drifting across block boundaries rather than resetting. This suggests participants continuously updated their beliefs about the drifting environment rather than settling on a fixed choice.

### EEG acquisition and preprocessing

EEG data were recorded at a sampling rate of 500Hz using a 64-channel ANT Neuro Waveguard Original cap using extended 10–20 montage (63 scalp Ag/AgCl electrodes and one EOG) connected to an eego mylab ANT amplifier and digitized at 24-bit resolution. The reference electrode was CPz and AFz was used as ground, with all the impedances kept below 10 kΩ. EEG recordings and stimulus presentations were synchronized using the LabStreamingLayer software Labrecorder (LSL Labrecorder version 1.13; Kothe et al., 2025). The data were preprocessed offline using the BeMoBIL Pipeline (Klug et al., 2022) with the EEGLAB toolbox (Delorme & Makeig, 2004; EEGLAB version 2026.0.0) following the procedure used in previous studies (Naveilhan et al., 2025, 2026; Saulay-Carret et al., 2026).

For the preprocessing we first downsampled the signal to 250 Hz offline, and after visual inspection of the raw signal, segments containing substantial artifacts and non-experimental periods were removed. Line noise was then removed using Zapline-plus (Klug & Kloosterman, 2022), which detects noise peaks automatically. Bad electrodes were identified using the BeMoBIL pipeline’s automated channel detection, which flags a channel as abnormal if its correlation with a robust estimate based on other channels falls below 0.8 and if it remains broken for more than 50% of the recording. This detection procedure was repeated ten times, and a channel was rejected if flagged in 50% or more of the repetitions. On average, 1.98 ± 0.98 channels per participant were rejected and interpolated using spherical spline interpolation. The number of channels interpolated did not differ between groups (control 2.04 ± 1.19 *vs.* ADHD 1.91 ± 0.67; Welch *t*_(42.0)_ = 0.46, *p* = .65). The data were then re-referenced to the common average of the 63 scalp channels (the EOG channel was excluded from the reference).

For the spatial decomposition, a copy of the data was high-pass filtered with a passband edge at 1.5 Hz and decomposed with AMICA (Klug & Gramann, 2021; Palmer et al., 2008), using one model and 2,000 iterations. Time periods were automatically rejected during the decomposition using AMICA’s built-in rejection procedure, which iteratively removes data samples exceeding 3 standard deviations from the mean over 10 rejection cycles, leading to an average of 3.43 ± 1.33% of rejected samples (without any difference between groups: control 3.30 ± 1.12% *vs.* ADHD 3.58 ± 1.54%; Welch *t*_(39.5)_ = +0.71, *p* = .48). Equivalent single-dipole models were fitted to each component with DipFit, discarding dipoles outside the head model, and components were labelled with ICLabel (Pion-Tonachini et al., 2019, *default* classifier). The labels were then copied to the unfiltered data, and we retained only the ones labelled as brain by the classifier, leading to an average retention of 12.68 ± 3.90 components per participant (range 6–21). The number of retained components did not differ between groups (control 12.78 ± 3.48 *vs.* ADHD 12.57 ± 4.43; Welch t_(41.5)_ = 0.19, *p* = .85).

The brain-component-reconstructed cleaned signal was then band-pass filtered between 0.3 and 40 Hz and epoched from −400 to +800 ms around the choice and payoff phases for each trial, with a baseline of −200 to 0 ms. The number of trials entering analysis was slightly lower in the ADHD group (control 443.5 ± 7.6 *vs.* ADHD 434.1 ± 12.8; Welch *t*(34.6) = −3.07, *p* = .004), because a greater amount of data were removed during manual artifact rejection. This difference is unlikely to affect the interpretation of our results, for two reasons. First, measurement precision did not differ between groups: the standardized measurement error (SME, Luck et al., 2021) of the parietal P300 at the decision phase was 0.143 µV in controls and 0.188 µV in the ADHD group, a difference that was not significant (Welch *t*_(25.7)_ = 1.18, *p* = .248; Mann– Whitney *p* = .508). Second, across participants, the SME was essentially uncorrelated with the number of trials contributing to each average (*r* = −0.15, *p* = .31), and a pure trial-count effect would predict an ADHD/control SME ratio of only 1.01, whereas the observed ratio was 1.31. Any residual difference in precision therefore reflects single-trial variability rather than the number of trials retained, ensuring that between-group electrophysiological differences are unlikely to be attributable to differential data loss or disparities in signal-to-noise ratio.

### Statistical analyses

All the behavioral statistical analyses were done using Python 3.11 (*NumPy*, *SciPy*, *statsmodels*, *scikit-learn*, *PyMC*) and R 4.5.1. Group comparisons for behavioral measures were conducted using two-tailed Welch’s *t*-tests, which do not assume equal group variances (Delacre et al., 2017). Effect sizes are reported as Hedges’ *g*, incorporating the standard small-sample correction for upward bias (Hedges, 1981). To align with the assumptions of Welch’s test, the mean difference was standardized using an unweighted average of group variances rather than weighting by sample size (Bonett, 2008); although both approaches yield virtually identical values given the sample size. Confidence intervals (95% CI) were calculated from the standard error of *g* using the Welch–Satterthwaite degrees of freedom (Lakens, 2013).

#### Behavioral analyses

### Behavioral measures

Performance was quantified as the percentage of the maximum available payoff, calculated by dividing the payoff obtained by the highest payoff available on each trial and averaging these ratios across trials, and as the percentage of objectively optimal choices, defined as selecting the option with the highest latent reward mean.

Choices were classified online using an empirical criterion that tracked the most recently observed payoff for each alternative prior to the current trial. An action was classified as exploitative if the participant selected the option with the highest last-observed payoff, and exploration otherwise. This operational definition relies directly on observable reward history rather than unobservable latent states or model-derived expected values, allowing trial labels to be generated in real time and recorded synchronously with EEG event markers. Control analyses confirmed that this heuristic classification had high concordance with model-based alternatives computed *a posteriori*: showing 94.3% agreement (Cohen’s κ = .86) with a mean- reverting Kalman filter observer (Daw et al., 2006) and 96.6% agreement (Cohen’s κ = .91) with a fitted dual-learning-rate reinforcement learning model (Lefebvre et al., 2017).

Crucially, exploration was dissociated from behavioral switching (defined as selecting a different option from the preceding trial). Under our criterion, switching to the option with the highest last-observed payoff constitutes an exploitative switch, whereas repeating an option with an inferior last-observed payoff constitutes exploratory sampling. Although the two classifications were moderately correlated (ϕ=.58), they captured distinct behavioral patterns: 37.7% of all switches were exploitative, and 20.1% of exploratory choices involved repeating the previous action.

### Reward-history kernel

The influence of recent outcomes on choice was first characterized using a single-predictor logistic regression fit separately for each participant, relating the payoff received on trial *t-1* (in points) to the probability of repeating that option on trial *t*. The resulting slope estimates were compared between ADHD and control groups. To complement this parametric analysis, choice repetition was also examined categorically by contrasting the probability of staying with the same option following a low payoff (below the individual median) versus a high payoff.

To capture the extended temporal profile of reward dependency, a multivariable logistic regression kernel was estimated (Lau & Glimcher, 2005). This model predicted choice repetition on trial *_t_* from the payoffs received in the previous six trials, with lags of 1 to 6 (*t-1* to *t-6*), with reward values z-scored within-participant so that regression weights reflect standardized effect sizes per standard deviation of reward while mutually adjusting for all other lags. Lag-specific weights were first evaluated across participants using two-tailed one-sample *t*-tests against zero to characterize the kernel’s overall decay shape. To test group differences in the lag-1 weight with maximal statistical efficiency, the model was refit at the single-trial level using a Generalized Estimating Equation (GEE) with an exchangeable working correlation matrix clustered on participant, specifying diagnostic group as a predictor of both the intercept and the lag-1 weight. Finally, the relationship between an individual’s lag-1 weight and overall task performance (percentage of maximum available points earned) was evaluated using Spearman rank correlations computed separately within each group, preventing spurious associations due to group-level pooling.

### Hierarchical Bayesian model of the reward kernel

To complement the GEE test with a continuous, directly interpretable measure of evidence, we re-estimated the lag 1–6 reward kernel within a hierarchical generalized linear model (Bernoulli likelihood, logit link). This notably allowed us to quantify evidence in favor of the absence of group differences at longer lags, an inference that non-significant frequentist *p*-values cannot provide. The model specified participant-level varying intercepts and varying slopes for the lag-1 reward (partial pooling), while diagnostic group was entered as a population-level offset on the intercept, on the primary lag-1 reward weight, and on the reward weights of lags 2 to 6. Priors were chosen to be weakly informative: "Normal" (0,2.5) on the population intercept, "Normal" (0,2.0) on the population lag-1 weight, "Half-Normal" (0,2.0) and "Half-Normal" (0,1.0) on the respective between- participant standard deviations, "Normal" (0,1.0) on the primary group offset, and "Normal" (0,0.5) on the offsets at lags 2 to 6. Posteriors were estimated using the No-U-Turn Sampler (NUTS; Hoffman & Gelman, 2014) across 4 chains of 1,500 sampling draws following 1,500 warmup/tuning iterations, with a target acceptance probability set to 0.92. Convergence was confirmed across all parameters (R-hat ≤ 1.01, minimum bulk effective sample size = 638). Parameter inference is reported via posterior means alongside 95% highest density intervals (HDIs), the directional posterior probability that the group offset exceeds zero (P_(Δβ>0)_), and Savage–Dickey Bayes factors evaluated at zero against the prior density to quantify evidence for the null hypothesis (BF_01_) versus the alternative (BF_10_; Wagenmakers et al., 2010).

## ERP analyses

### Cluster-based permutation

Following Hassall et al. (2019), EEG analyses targeted two predefined regions of interest: midfrontal (FCz, FC1, FC2, Fz, F1, F2) and centro-parietal (Pz, P1, P2, CP1, CP2) clusters. Condition and group effects were evaluated across the entire -200 to +800ms epoch for each event (decision and payoff phases) using non-parametric cluster- based permutation testing (Maris & Oostenveld, 2007).

Within each group, condition differences (explore vs. exploit; high vs. low payoff) were tested against zero at every time sample using two-tailed paired *t*-tests, with empirical null distributions constructed via random sign-flipping of participant-level difference waveforms (5000 permutations). Between-group differences were evaluated by contrasting these condition- difference waveforms across groups at every time point using two-sample *t*-tests, with null distributions generated by randomly permuting group labels (5000 permutations). In both analyses, temporally contiguous samples exceeding an uncorrected cluster-forming threshold of |t| ≥ 2.0 were grouped into candidate clusters. The test statistic for each cluster was defined as the sum of its constituent *t*-values (cluster mass or "maxsum"). A cluster was deemed statistically significant if its mass exceeded the 97.5th percentile of the maximum cluster-mass distribution across the 5000 permutations (two-tailed α = .05). The identical cluster-forming threshold and cluster-mass statistic were applied across all electrophysiological analyses.

Because cluster-based tests can be too conservative for effects that are time-locked or consist of a latency shift, the two component effects (P3 in the decision phase, centro-parietal region of interest [ROI]; P2 in the payoff phase, midfrontal ROI) were further quantified in windows defined on a grand average collapsed across groups and conditions, which is orthogonal to both contrasts (Kriegeskorte et al., 2009; Luck & Gaspelin, 2017). The window was set at ±50 ms around the collapsed-average peak and applied identically to all participants; amplitude was the mean voltage over the window and ROI, latency the 50% fractional area latency.

Because exploratory trials were less frequent, we also repeated this analysis after subsampling exploitation trials within-participant to control for trial counts. In both the full and matched analyses, no group (ADHD/Control) × condition (Explore/Exploit) cluster reached significance for any of the 2 ROIs (smallest cluster *p* = .175 unmatched, .200 matched).

### Single-trial analysis of outcome change

To examine how electrophysiological responses varied with changes in recently received rewards, signed outcome change was calculated as the payoff received on the current trial minus the payoff received on the preceding trial. Unsigned outcome change was defined as the absolute value of this difference and was compared between groups using cluster-based permutation tests with 5,000 permutations. The unsigned difference averaged over the identified cluster was subsequently used as the outcome variable in the exploratory symptom-prediction analysis. To examine the frequency content of this response, the same analysis was repeated in an exploratory manner on signals filtered into the 0.3–2, 2–4 (delta), and 4–8 Hz (theta) bands and on a version high-pass filtered above 2 Hz.

## Results

### No evidence for greater exploration or altered explore/exploit ERPs in ADHD

First, we aimed to test our primary hypothesis that ADHD participants would explore more compared with the control group, leading to less optimal performance, as previously reported. However, we detected no statistically significant group differences in task performance (**Figure 2.a** and **b**): the two groups obtained a similar percentage of the maximum gain available on the trials they completed (control 91.7 ± 2.5%; ADHD 92.7 ± 2.0 %; *t*_(47.8)_ = 1.58, *p* = .121, *g* = 0.43) and a similar proportion of objectively optimal choices (63.1% vs 65.9%; *t*_(45.6)_ = +1.19, *p* = .239, *g* = 0.32). Reaction times were likewise indistinguishable (mean 498 vs 499 ms; *t*_(44.5)_ = 0.04, *p* = .971, *g* = 0.01), as was reaction-time variability (CV 0.435 vs 0.465; *t*_(43.8)_ = 1.06, *p* = .294, *g* = 0.30). We also found no significant evidence of greater exploration in the ADHD group: exploratory choices represented 24.9 ± 7.1 % of trials compared with 28.0 ± 10.2 % in controls, with a non-significant difference in the direction opposite to the prediction (*t*_(46.4)_ =−1.28, *p* = .206, *g* = −0.35), and the raw switch rate showed the same pattern (33.0% vs 35.0%; *t*_(48)_ = −0.57 *p* = .570, g = -0.16).

**Figure 2.**
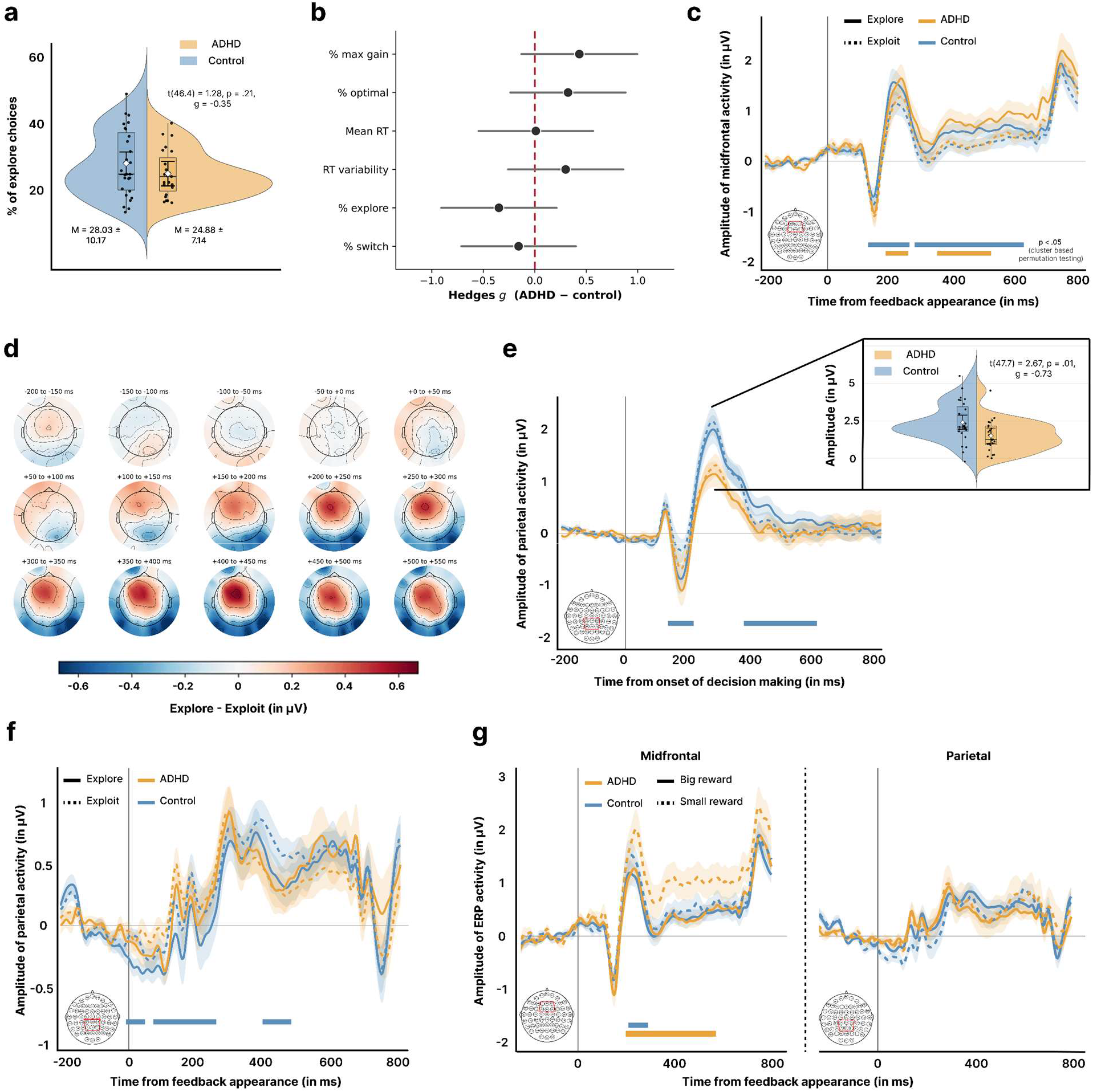
Presentation of the results. **(a)** Percentage of exploratory choices per participant (violin plots with individual data points); groups did not differ significantly. **(b)** Effect sizes (Hedges’ g, ADHD − control) with 95% CIs for behavioral measures: % of maximum gain, % optimal choices, mean RT, RT variability, % explore, and % switch; none reached significance. **(c)** Midfrontal ERP amplitude (FCz, FC1, FC2, Fz, F1, F2) time-locked to feedback onset, split by explore (solid) vs. exploit (dashed) trials and by group; horizontal bars indicate significant cluster-based permutation testing, with no group × condition interaction. **(d)** Scalp topographies of the explore–exploit difference across successive 50 ms windows post-feedback, showing a midfrontal-focused effect. **(e)** Parietal ERP amplitude (CP1, CP2, P1, Pz, P2) time-locked to decision onset, split by explore vs. exploit trials; inset shows the P3 peak amplitude reduced in ADHD relative to controls. **(f)** Parietal ERP amplitude time-locked to feedback onset, split by explore vs. exploit and group; significant clusters were present for controls only. **(g)** Midfrontal (left) and parietal (right) ERP amplitude time-locked to feedback onset, split by reward magnitude (big vs. small reward) and group; a significant cluster emerged in midfrontal electrodes, larger in ADHD than controls, and absent parietally.

Regarding the ERPs, we found a modulation of midfrontal and centro-parietal activity when contrasting explore and exploit trials, in line with previous work. In the midfrontal electrodes (FCz, FC1, FC2, Fz, F1, F2), we found an increased activity for the explore trials for both groups. Cluster-based permutation testing indicated that this difference was present for a first cluster around 200ms, and a second one lasting up to the end of the feedback phase without any group interaction for any of the clusters (**Figure 2.c**). This effect was mainly localized to these electrodes, as shown in **Figure 2.d,** although some differences were also present for the centro- parietal region of interest (CP1, CP2, P1, Pz, P2), with 3 clusters of significant difference for the control group only (**Figure 2.f**). However, we found no interaction between group and the explore-exploit choice for either of the regions of interest in the feedback or decision-making phases (smallest cluster *p* = .200).

The only difference observed between ADHD and control participants concerned the P3 elicited in the decision phase, over the centro-parietal electrodes, which was reduced in amplitude (2.41 vs 1.51 µV; *t*_(47.7)_ = −2.67, *p* = .010, *g* = −0.73, **Figure 2.e**). This difference was independent of the explore/exploit distinction, as the P3 was not itself modulated by that contrast in either group (*p* = .31), and the corresponding group × condition interaction was null (all *p* ≥ .28).

In a second analysis to test our hypothesis that the ADHD group should differ in reward processing, we compared the trials where the payoff was a low (below the median) with those where it was high, irrespective of the explorative nature of the trial. This was motivated by the prediction-error literature, which suggests that the amplitude of feedback-related potentials scales with the magnitude of the outcome rather than reflecting a purely binary win/loss signal (Holroyd & Coles, 2002). Here, we found that adults with ADHD displayed a strong increase in midfrontal ERP amplitude for small compared to big rewards within a cluster starting around 200 ms and lasting until the end of the feedback period, a pattern that was also present but less pronounced in controls, and absent over parietal electrodes **(Figure 2.g**). Analysis of the midfrontal P2 peak amplitude confirmed this heightened sensitivity in ADHD, showing a significantly larger difference between small and big rewards compared to controls (+ 0.996 µV comparing the small and the big rewards for ADHD vs + 0.332 µV for control, *t*_(38.34)_ = 2.47, *p* = .018, *g* = 0.71 [0.13, 1.29]). Thus, although groups did not differ in overall explore–exploit behavior, these findings suggest that participants with ADHD may have solved the task through distinct strategies driven by differential reward weighting, which motivated the subsequent analyses.

### Recent rewards exert a stronger influence on choice and midfrontal outcome- change responses in ADHD

Although no significant group differences were detected in exploration frequency or overall performance, the planned reward-history analyses indicated a stronger association between the most recent reward and subsequent choice repetition in the ADHD group. This was first observed when comparing the probability of staying on the current option according to the payoff received in the previous trial (**Figure 3.a**). Fitting a logistic function linking the payoff received to the probability of staying on the same option, the slope was steeper in ADHD than in control participants (1.50 vs 1.14 per 10 payoff points; t_(43.6)_ = 2.80, *p* = .008, g = 0.79, 95% CI [0.21, 1.37]). After a low payoff (below median) ADHD participants stayed on 12.1% of trials against 19.1% in controls, t_(46.9)_ = −2.28, *p* = .027, g = −0.64, 95% CI [−1.21, −0.07]. Conversely, after a high payoff (above median) ADHD stayed on 92.9% against 87.9% (t_(47.9)_ = +2.14, *p* = .037, g = +0.59, 95% CI [+0.02, +1.16]). Thus, rather than displaying an indiscriminate tendency to explore or exploit, participants with ADHD exhibited a more pronounced win-stay / lose-shift policy, characterized by sharper behavioral contingency on the immediate outcome.

**Figure 3.**
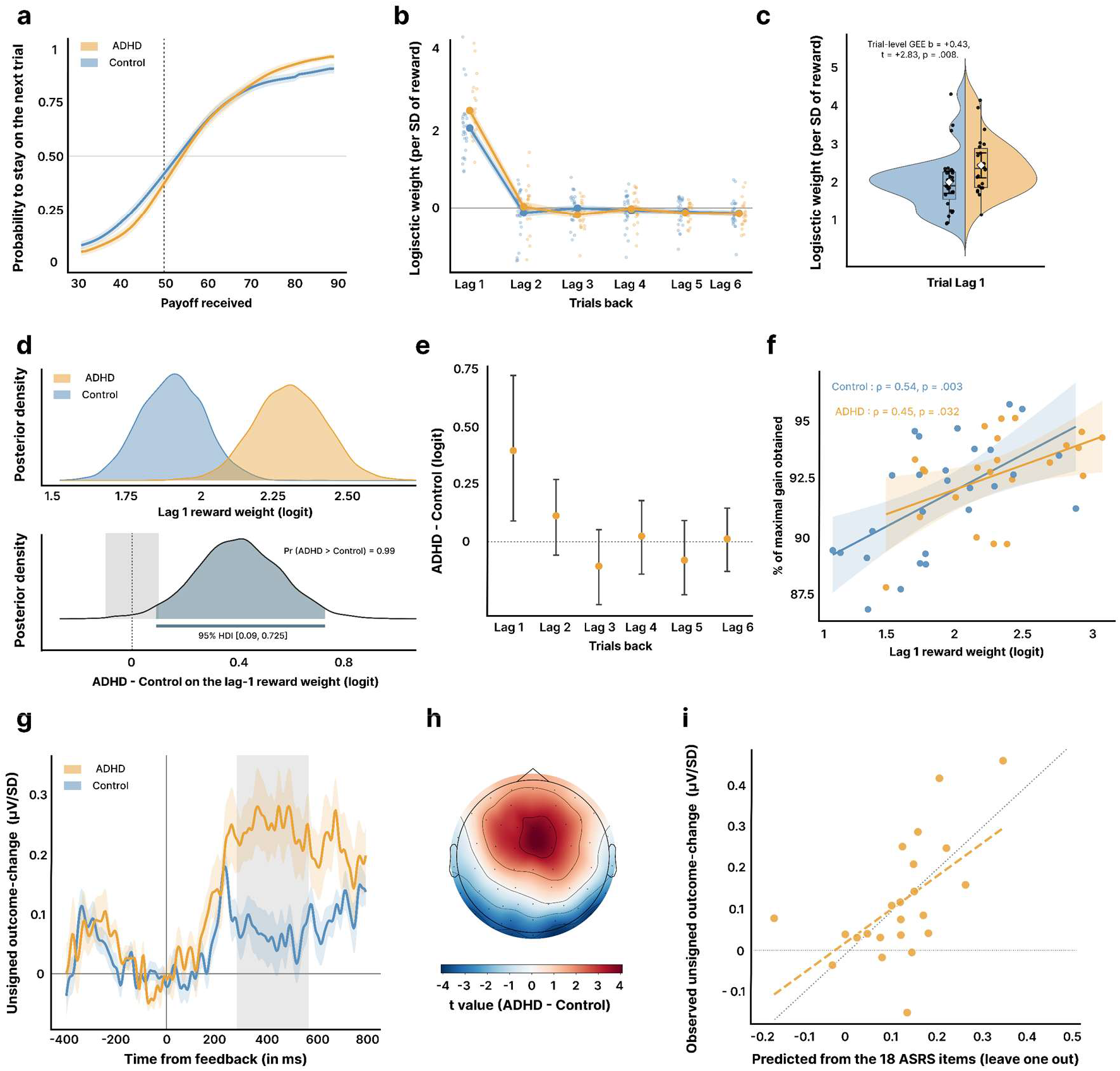
Reward-history effects and midfrontal sensitivity to unsigned outcome change. **(a)** Probability of staying on the same option as a function of the payoff received on the previous trial, fitted with a logistic function per group; the slope was steeper in ADHD than in controls. **(b)** Logistic weights (per SD of reward) for rewards received one to six trials back, mutually adjusted; the lag-1 weight was large and positive in both groups, while earlier lags carried small negative weights. **(c)** Lag-1 weight per participant (violin plots with individual data points), larger in ADHD as reported by the trial-level GEE group × lag-1 reward interaction. **(d)** Hierarchical Bayesian model: posterior densities of the lag-1 reward weight per group (top) and of the ADHD − control difference with its 95% HDI (bottom), which excludes zero. **(e)** Bayesian group offsets (ADHD − control) with 95% HDIs at each lag from 1 to 6; only lag 1 excludes zero. **(f)** Association between the lag-1 reward weight and the percentage of maximum available gain obtained, positive in both groups. **(g)** Feedback-locked midfrontal regression coefficients for unsigned outcome change, shown separately by group. Coefficients quantify the change in ERP amplitude associated with a one-standard-deviation increase in the absolute difference between consecutive payoffs, controlling for signed outcome change. The shaded interval indicates the significant group-difference cluster (280–568 ms). **(h)** Scalp distribution of the group contrast in unsigned outcome-change coefficients, expressed as t values averaged over the cluster interval. **(i)** Observed versus cross-validated predicted unsigned outcome-change coefficients within the ADHD group, using the 18 ASRS item scores as predictors in a ridge regression.

To determine whether this heightened outcome sensitivity was confined strictly to the most recent outcome or reflected an altered temporal integration window, we modeled choice repetition across an extended reward kernel spanning lags 1 to 6 (**Figure 3.b**). Payoffs were standardized (*_z_*-scored) within participants, yielding mutually adjusted regression weights per standard deviation of reward. Across both groups, choices were governed predominantly by the most recent trial: the lag-1 weight was large and positive (t_(49)_ = 20.32, *p* < .001), whereas earlier lags each exerted a small negative influence (lag 2 −0.12 to lag 6 −0.36, all *p* < .05 against zero). In a trial-level GEE clustered on participant, the reward received on the previous trial strongly predicted staying (b = 1.82, *p* < .001), and this effect was larger in ADHD participants (group × lag-1 reward: b = 0.43, 95% CI [0.15, 0.73], *p* = .008, **Figure 3.b** and **c**).

This group difference was supported by a hierarchical Bayesian model of the same kernel (trial- level Bernoulli likelihood, partial pooling, random slopes on the lag-1 reward), which estimated a population-level ADHD offset of +0.397 logit units, with a 95% HDI of [+0.090, +0.725] excluding zero (P_(ADHD−Control > 0)_ = 0.990), and moderate evidence for a group difference (Savage–Dickey BF_10_ = 3.12; Figure 3.d). Because all lag predictors were estimated simultaneously within the multivariable kernel, these weights reflect mutually adjusted partial effects. For lags 2 through 6, the 95% HDIs of the group offsets all included zero ([−0.06, +0.27], [−0.28, +0.05], [−0.14, +0.18], [−0.23, +0.09], [−0.13, +0.15]; BF_01_ = 5.7, 4.9, 12.4, 8.9 and 14.5 for lags 2 to 6, respectively; Figure 3.e), providing moderate (lags 2 and 3) to strong (lags 4 to 6) evidence against between-group differences in the weighting of outcomes beyond the immediately preceding trial. It is important to note that in this environment the strategy was not maladaptive, and the lag-1 weight correlated with the percentage of the maximum available gain obtained for both ADHD (ρ = 0.45, *p* = .032) and controls (ρ = 0.54, *p* = .003), suggesting that prioritizing the most recent outcome facilitated overall performance (**Figure 3.f**).

At the neural level, feedback-locked midfrontal ERP amplitude varied more strongly with unsigned outcome change in the ADHD group than in controls. The group difference formed a significant cluster spanning 280–568 ms after feedback onset (cluster-based permutation *p* = .002; **Figure 3.g**), with a predominantly midfrontal scalp distribution (**Figure 3.h**). This association reflected the magnitude of the difference between consecutive payoffs, with signed outcome change included simultaneously in the regression. Within the ADHD group, an exploratory ridge regression examined whether the combined scores of the 18 ASRS items predicted the unsigned outcome-change coefficient averaged over this cluster. With the regularization penalty selected through cross-validation within each training fold, leave-one- participant-out predictions yielded Q² = .303 and a correlation between observed and predicted coefficients of r = .573 (permutation *p* = .010; **Figure 3.i**). We also performed correlations between the 3 dimensions of the ASRS and the unsigned outcome change, with a tendency only for the impulsivity dimension (see **Supplementary 1**).

The group difference in sensitivity to unsigned outcome change was predominantly associated with slow ERP activity. In the 0.3–2 Hz signal, a significant cluster spanned 280–568 ms after feedback onset (cluster *p* = .018). No significant clusters were identified in the 2–4 Hz or 4–8 Hz signals (smaller *p* = .34 and *p* = .44, respectively). High-pass filtering above 2 Hz reduced the estimated group difference by 91%, and the remaining difference was not statistically significant (*p* = .67). Together, these findings indicate that the observed difference was primarily expressed in the slow component of the evoked response, below 2Hz.

## Discussion

The present study examined whether adults with a diagnosis of ADHD explore more frequently and show greater neural differentiation between exploration and exploitation, alongside stronger responsiveness to reward in comparison with neurotypical participants. Neither behavioral exploration rates nor neural exploration–exploitation ERP contrasts differed significantly between groups. However, ADHD participants showed a stronger reliance of subsequent choices on the most recent reward and greater midfrontal ERP sensitivity to unsigned changes between consecutive outcomes. These differences occurred without significantly poorer performance, suggesting that ADHD-related differences concerned responsiveness to recent feedback rather than an overall increase in exploration *perse*.

Contrary to our first hypothesis, the present study did not detect significantly greater exploration in adults with ADHD. This discrepancy with previous observations may reflect differences in task demands, latent choice processes, and sample characteristics. In the present task, exploiting recently rewarding options was advantageous, as reflected in the positive association between recent reward weighting and earnings across both groups. While exploration remained necessary to track drifting alternatives, the reward structure likely encouraged sustained exploitation. This is consistent with evidence that exploratory behavior depends on the context in which the task is embedded. For example, Addicott et al. (2021) reported greater exploration and lower earnings in clinically diagnosed adults with ADHD performing a six-armed bandit task, whereas Barack et al. (2024) found earlier patch departure and higher reward rates in an online foraging task. Second, aggregate exploration rates may obscure differences in underlying generative processes. Directed exploration (seeking information) and random exploration (choice variability) respond differently to uncertainty and future reward opportunities (Gershman, 2018; Wilson et al., 2014). Although our payoff-based classification agreed closely with Kalman-filter and fitted-value models (94.3% and 96.6%, respectively), it does not establish whether exploratory choices reflected information-seeking or stochasticity. The nonsignificant group difference therefore concerns exploration frequency under our operational definition, without ruling out divergent latent mechanisms. Finally, our sample size and our recruitment of university students limit generalization across the broader adult ADHD spectrum (Faraone et al., 2024). Individuals pursuing higher education often possess stronger compensatory scaffolding (DuPaul et al., 2009; Weyandt et al., 2013), potentially buffering decision-making vulnerabilities that would otherwise emerge in less structured contexts. Nevertheless, the ADHD group reported elevated ASRS scores and showed the attenuated decision-locked P300 commonly reported in adult ADHD (Kaiser et al., 2020; Szuromi et al., 2011), suggesting that this sample shares key features of clinical ADHD samples.

Despite equivalent overall exploration rates, groups differed in reward sensitivity, and ADHD participants were more likely to repeat choices following high rewards and abandon them following low rewards. These opposing tendencies can balance across trials, leaving aggregate exploration and switching rates relatively unchanged. Reward-history analyses localized this group difference to the immediately preceding outcome, with no evidence for differences at earlier lags. This partly supports the predicted influence of high rewards while extending it to greater responsiveness to low rewards. Such a pattern is more specific than generalized reward- seeking and fits accounts emphasizing altered reinforcement contingencies and the temporal weighting of outcomes in ADHD (Luman et al., 2005, 2010; Tripp & Wickens, 2008). Within multiple-pathway accounts of ADHD (Sonuga-Barke et al., 2010; Sonuga-Barke, 2002), it points to the motivational, reward-related pathway rather than to executive dysfunction, consistent with the absence of group differences in overall performance and response times. Interestingly, stronger recent reward weighting was positively associated with earnings within both groups. In uncertain contexts, recent observations are informative, though their usefulness depends on balancing genuine environmental change against observation noise (Behrens et al., 2007; Nassar et al., 2012). Greater responsiveness may facilitate adaptation when values change, despite risking unnecessary switches after unrepresentative outcomes. While not establishing causality, this association challenges an inherently maladaptive interpretation and is in line with evidence that ADHD decision-making difficulties do not only reflect greater risk- seeking (Dekkers et al., 2021). Clinically, a stronger reliance on the most recent outcome may relate to the tendency of adults with ADHD to change course after a single experience (negative or positive), and is consistent with the emphasis on frequent, immediate feedback in psychosocial interventions.

Beyond these results, a central neurocomputational question is whether this steeper stay–switch pattern reflects heightened reward sensitivity or reduced decision noise. Standard neurocomputational psychiatry accounts propose that ADHD involves blunted cortical neural gain and elevated decision noise, predicting increased random exploration (Dubois & Hauser, 2022; Hauser et al., 2016). In contrast, our participants with ADHD displayed steeper stay slopes, non-significantly fewer exploratory choices, and slightly higher earnings, patterns consistent with a more deterministic, greedier choice policy. This apparent divergence can be explained by the task environment. In a restless bandit characterized by high volatility (high Kalman gain), recent payoffs closely approximate true current values, making past history negligible and rendering a simple win-stay/lose-shift heuristic near-optimal. Under these conditions, recent and salient rewards provide a strong exogenous signal that temporarily overrides elevated baseline cortical noise (Chakroun et al., 2020; Tripp & Wickens, 2008). One possibility is that choice stochasticity in ADHD is dynamically modulated by reward recency.

Paralleling behavioral results, neural differences reflected responsiveness to recent outcomes rather than overall exploration frequency. Previous studies have reported enhanced feedback processing around exploratory decisions (Hassall et al., 2019; Hassall & Krigolson, 2020), but we did not detect significantly greater exploration–exploitation ERP differentiation in ADHD. Instead, group differences emerged in the association between feedback-evoked amplitude and unsigned outcome change. This suggests that sensitivity to recent outcome variations captures differences that are obscured when trials are classified solely as exploratory or exploitative. Specifically, adults with ADHD showed a stronger midfrontal association with unsigned outcome change within a cluster extending from 280 to 568 ms after feedback onset. Because signed and unsigned changes were entered simultaneously, this association captured sensitivity to the magnitude of consecutive payoff differences while controlling for outcome direction. This finding indicates greater neural responsiveness to feedback shifts, complementing the stronger behavioral weighting of the preceding reward. To test whether this reflected prediction- error scaling rather than heuristic comparisons, we evaluated trial-by-trial prediction errors from computational reinforcement-learning models (**Supplementary 2**) and found that midfrontal voltage scaled more strongly with prediction-error magnitude in ADHD across both positive and negative outcomes.

To better understand this result we then investigated the spectral profile of this ERP’s midfrontal response, and found that the group difference was driven primarily by slow activity below 2 Hz rather than faster theta rhythms (4–8 Hz) that typically index cognitive conflict or initial prediction errors (Cavanagh & Frank, 2014). Under the classical “context-updating” framework, frontocentral slow waves are proposed to reflect how the brain revises its internal model when incoming feedback signals an environmental change (Donchin, 1981; Polich, 2007). In a drifting task where sudden payoff shifts indicate changing contingencies, this heightened slow-wave sensitivity suggests that adults with ADHD update their internal representation of the context more strongly after salient outcome changes (*i.e*., magnitude of the positive or negative outcome). While medial frontal circuits are central to feedback evaluation and behavioral adjustment (Blanchard & Gershman, 2018; Frank et al., 2015; Hauser, Iannaccone, Ball, et al., 2014), and align with prior neuroimaging evidence linking ADHD decision-making to frontostriatal dysfunction, exploratory tendencies, and altered reward-modulated control (Jones et al., 2020; Norman et al., 2018), our data propose that feedback reactivity in ADHD can be interpreted as an enhanced late model-updating response rather than an early conflict signal. More broadly, this explanation converges with predictive- coding accounts proposing that ADHD is characterized by a disproportionate weighting of incoming evidence relative to prior expectations (Gonzalez-Gadea et al., 2015). Consistent with this view, elevated ADHD symptoms have recently been associated with a diminished influence of perceptual history on current judgments (Marsicano et al., 2026) and attentional lapses may impair the incorporation of prior knowledge into perceptual decisions (Morales et al., 2015). In our study, this translates to over-weighting the most recent payoff relative to the accumulated reward statistics of each option, a pattern directly reflected in the reported selective group difference at lag 1 and the absence of differences at longer lags. Importantly, in our drifting bandit task, empirical learning rates approached unity, expected values tracked outcomes rapidly, and stay-trial prediction errors correlated strongly with consecutive payoff differences. Future paradigms featuring lower environmental volatility will therefore be essential to corroborate these mechanisms.

Several methodological limitations of the present study should also be considered. First, our sample was powered for large effects on the primary behavioral measure (exploration rate), but it remains relatively modest. Consequently, the study may have lacked the sensitivity to detect subtler group differences or to conduct well-powered subgroup analyses. Moreover, multiple- pathway models predict that only a subset of individuals with ADHD is affected along each pathway (Sonuga-Barke et al., 2010), and such heterogeneity may have attenuated group-level differences. Second, diagnostic classification relied on self-reported prior clinical diagnoses corroborated by the ASRS-v1.1 screening scale, rather than an in-person, semi-structured clinical interview. Finally, as we recruited predominantly from a university setting, our participants may have possessed greater compensatory scaffolding and cognitive reserve than the broader adult ADHD population, potentially masking some decision-making impairments that might emerge in less structured environments.

To conclude, contrary to our first prediction, we found no evidence that adults with ADHD explored more, switched more often, or showed greater midfrontal ERP differentiation between exploratory and exploitative choices. In contrast, our second hypothesis was supported and extended, as choices in ADHD relied more strongly on the immediately preceding outcome, marked by more frequent repetition after high rewards and more frequent switching after low rewards. At the neural level, this behavior was paired with an amplified midfrontal response to outcome changes, carried by slow activity below 2 Hz. Together, these results suggest that adults with ADHD weight newly incoming evidence more heavily than the reward history accumulated for each option, a result that can be interpreted in light of the predictive-coding framework. Overall, our results contribute to challenging purely deficit-centric models of ADHD, underscoring that its cognitive signature is not an invariant impairment, but a distinct mode of environmental engagement whose adaptive value can be shaped by contextual demands.

## Author Contributions

**Clément Naveilhan:** Conceptualization; Formal analysis; Investigation; Methodology; Writing-Original draft; Writing-Review & editing. **Remi Konan:** Conceptualization; Investigation. **Meggy Hayotte:** Writing-Review & editing; Supervision; Funding Acquisition. **Hervé Caci**: Writing-Review & editing. **Fabienne d’Arripe-Longueville:** Writing-Review & editing; Supervision; Funding Acquisition. **Stephen Ramanoel:** Conceptualization; Funding Acquisition; Methodology; Project administration; Supervision, Writing-Review & editing.

## Acknowledgements

The authors thank all participants whose time and commitment made this research possible. We also thank Dr. Valeria Manera, Dr. Lyne Daumas, Dr. Pierre Crescenzo and la Mission Handicap Université Côte d’Azur for insightful comments on the project and recruitment of the participants. This work has been supported by the French government, through the UCA^JEDI^ Investments in the Future project managed by the National Research Agency (ANR) with the reference number ANR-15-IDEX-01.

## Data and code availability

All raw and processed data, task materials, and custom analysis code supporting the findings of this study are publicly accessible on the Open Science Framework (OSF): https://osf.io/mjhn7/overview?view_only=e45ef603b8cf4fd18408ea20d41dc427

## Supplementary materials

### Supplementary 1

In this supplementary analysis, we decomposed the ASRS scores into three subdimensions (Inattention, Hyperactivity, and Impulsivity) and evaluated their relations with the midfrontal unsigned outcome-change modulation using both Pearson and Spearman correlations. All reported *_p_*-values are uncorrected for multiple comparisons. Across these analyses, only the impulsivity dimension exhibited a marginal, trend-level association with neural outcome sensitivity. Specifically, participants with ADHD who reported higher levels of impulsivity displayed steeper midfrontal regression slopes to unsigned outcome shifts. Neither inattentive nor hyperactive symptom scores showed a comparable relationship.

This selective trend between impulsivity and midfrontal outcome sensitivity provides an interesting nuance to our primary findings. In neurocomputational frameworks, trait impulsivity is closely tied to heightened reactivity toward immediate reward salience and elevated behavioral switching (Dubois & Hauser, 2022; Frank et al., 2015). Rather than indexing generalized inattention, the midfrontal slow-wave response to unsigned outcome change is proposed to reflect the dynamic context-updating and surprise tracking following salient payoff shifts. Within this framework, individuals scoring higher in impulsivity may exhibit an amplified neural "alerting" or model-updating signal when consecutive outcomes diverge sharply. This amplified midfrontal tracking of payoff volatility may serve as the neural driver of the steep win-stay/lose-shift policy identified behaviorally, prompting rapid behavioral reorientation whenever feedback deviates from expectations.

Although this association was exploratory and marginal after considering multiple comparisons, it aligns with evidence that the cognitive and electrophysiological heterogeneity of ADHD is organized along distinct symptom dimensions (Kaiser et al., 2020; Wilson et al., 2014). Future studies employing larger dimensional samples and structured clinical interviews will be valuable to confirm these results.

### Supplementary 2

To separate cumulative value updating from transient reward-driven repetition biases, we fitted computational reinforcement learning models combining valence-dependent learning rates with choice perseveration (Lefebvre et al., 2017). Payoffs were rescaled to 0–1, and option values were initialized at 0.5. Values were updated via a delta rule with separate learning rates for positive (α+) and negative (α−) prediction errors. A softmax choice rule integrated learned values with an outcome-independent perseveration parameter ("stickiness"; Katahira, 2015) and, in the full model, a reward-dependent repetition term scaled by the prior payoff centered at 0.5. Both repetition terms were set to zero on the first trial of each block. Maximum likelihood estimation (L-BFGS-B with eight random restarts) favored the full five-parameter model over a four-parameter model without reward-dependent repetition in 34 of 50 participants (mean BIC improvement of 15.5 points, median of 8.8 points). Parameter and model recovery confirmed high reliability for inverse temperature and repetition parameters (r = .94 to .96), with the full model successfully identified in 66% of synthetic simulations (33 of 50) and the simpler model in 100% (Wilson & Collins, 2019).

The fitted reward-dependent repetition weight was positive in 48 of 50 participants and significantly stronger in ADHD than in controls (7.98 vs. 5.27; t_(46.7)_ = 2.09, *p* = .042, g = 0.58), corroborating our model-free lag-1 findings. In exploratory analyses, learning rates exhibited valence asymmetry: controls updated values less after positive than negative prediction errors (α+ = .768, α− = .883; t_(26)_ = −2.28, *p* = .031), whereas adults with ADHD showed no reliable asymmetry (α+ = .904, α− = .846; t_(22)_ = 1.52, *p* = .143). The resulting between-group difference in asymmetry was significant (t_(46.4)_ = 2.73, *p* = .009, g = 0.74) and selectively driven by elevated positive learning rates in ADHD (*p* = .025; negative learning rates: *p* = .463).

However, this asymmetry finding warrants substantial caution. The effect was model- dependent, failing to reach significance in the reduced model omitting reward-dependent repetition (*p* = .082). Recovery precision was only moderate for learning rates (α+ r = .63, α− r = .45, asymmetry r = .48), statistical significance was reproduced in only one of three simulation replicates, and individual estimates frequently saturated at the 1.0 bound (21 for α+, 22 for α−). While such bound saturation is ecologically expected under rapid updating in non-stationary environments (Behrens et al., 2007), it may also denote trade-offs during optimization. Notably, recovery errors for α+ and reward-dependent repetition were uncorrelated (r = −.06, *p* = .45), and learning-rate asymmetry did not correlate significantly with empirical recent-reward metrics (r = −.001 to .140, all *p* >= .33), which themselves showed strong mutual convergence (r = .72 to .85).

Finally, we extended these results to the neural level and evaluated whether feedback-evoked midfrontal activity exhibited a valence-dependent group difference corresponding to the behavioral learning-rate asymmetry. Single-trial midfrontal voltage was regressed on model- derived prediction-error magnitude separately for positive (better than expected) and negative (worse than expected) outcomes, with slopes expressed on a common standardized scale (**Supplementary Figure 2**). Adults with ADHD exhibited steeper slopes across both valences. Significant group-difference clusters emerged between 252 and 332 ms for positive outcomes (+0.310 vs. +0.045 µV/SD, cluster *p* = .0395) and between 416 and 572 ms for negative outcomes (−0.486 vs. −0.134 µV/SD, cluster *p* = .0095). Because prediction errors of opposite valence elicit voltage deflections of opposite sign, steeper slopes on both sides indicate amplified midfrontal scaling to outcome magnitude generally, rather than a valence-selective counterpart to positive learning rates. Furthermore, because elevated learning rates induced strong correlation between modeled prediction errors and consecutive-payoff differences on stay trials, this analysis serves to demonstrate bilateral outcome sensitivity across valences rather than providing an orthogonal computational dissociation.

**Supplementary Figure 1.**
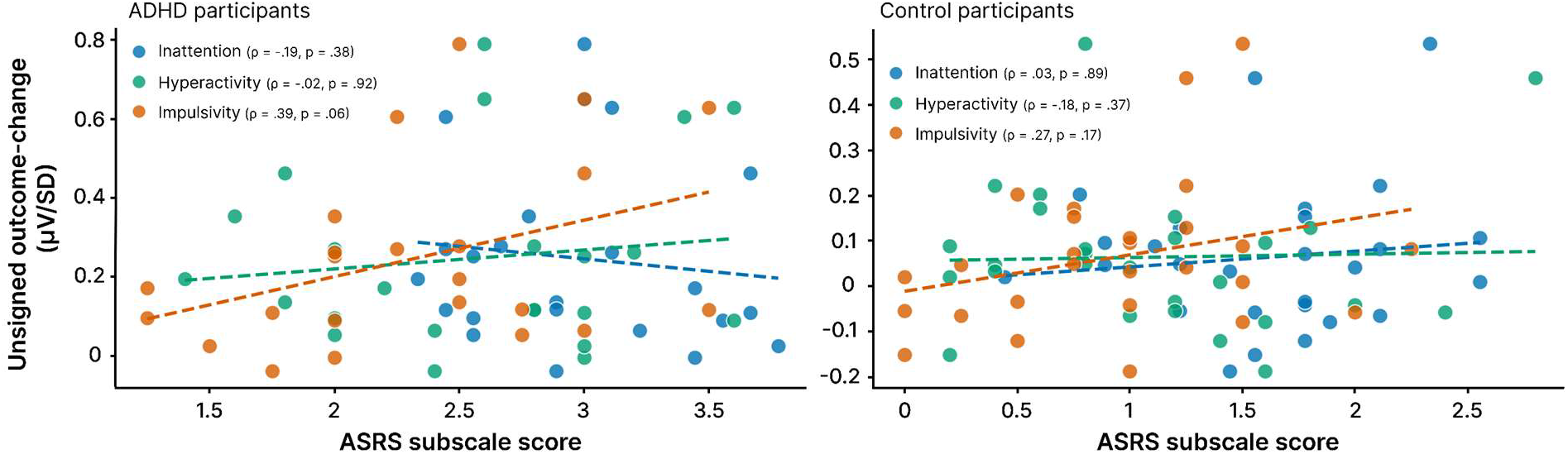
Correlation between ASRS subscales and neural outcome changes across groups. Scatter plots displaying unsigned outcome-change (µV/SD) against ASRS subscale scores in ADHD (left; n = 23) and Control (right; n = 27) groups. Data points and dashed regression lines represent Inattention (blue), Hyperactivity (green), and Impulsivity (orange) mean score. In the ADHD cohort, correlations were: Inattention (Pearson r = -0.12, *p* = .58; Spearman ρ = -0.19, *p* = .38), Hyperactivity (r = 0.13, *p* = .55; ρ = -0.02, *p* = .92), and Impulsivity (r = 0.39, *p* = 0.06; ρ = 0.39, *p* = 0.06). In the Control cohort, correlations were: Inattention (r = 0.11, *p* = .57; ρ = 0.03, *p* = 0.89), Hyperactivity (r = 0.03, *p* = .88; ρ = -0.18, *p* = .37), and Impulsivity (r = 0.29, *p* = .14; ρ = 0.27, *p* = .17).

**Supplementary Figure 2.**
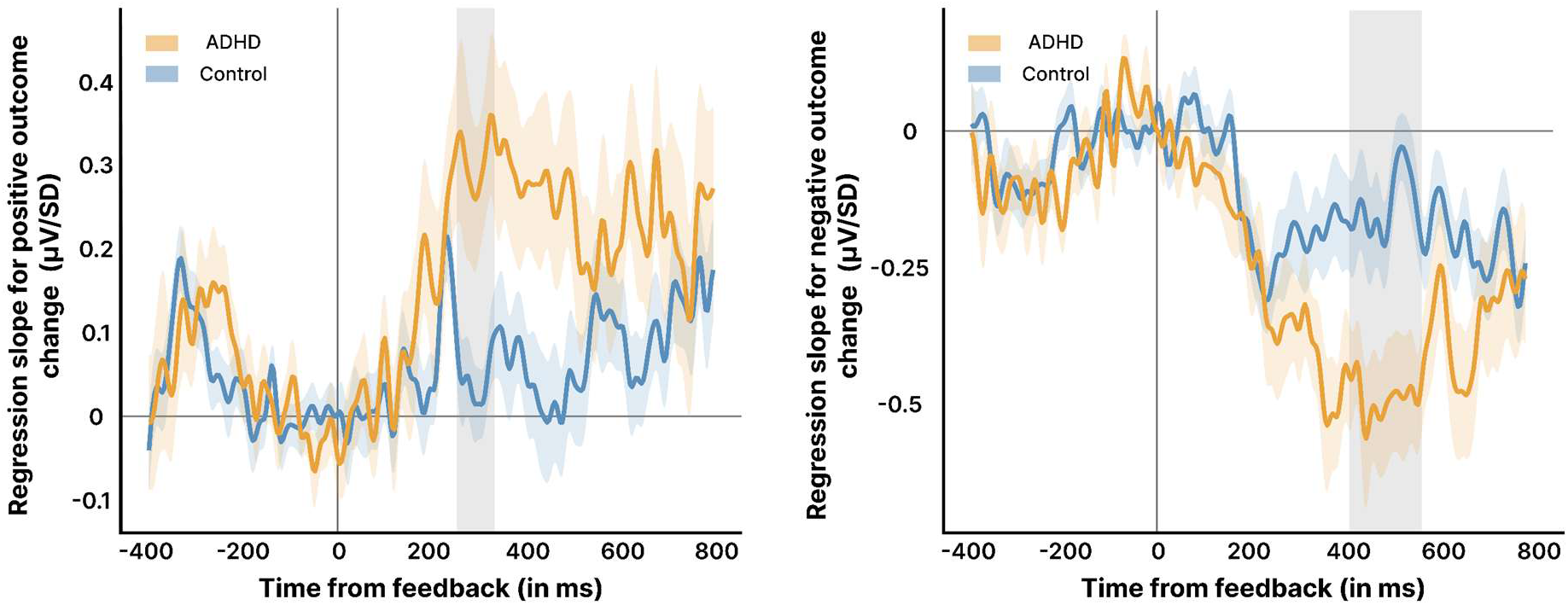
Midfrontal sensitivity partitioned by outcome valence (positive vs. negative prediction errors). Single-trial midfrontal voltage was regressed on model-derived prediction-error magnitude separately for positive (better than expected, left) and negative (worse than expected, right) outcomes, standardized as µV/SD. Solid lines denote group means (orange: ADHD; blue: controls), shaded ribbons indicate SEM, and the vertical line marks feedback onset. Shaded grey bands indicate significant between-group permutation clusters: 252–332 ms for positive outcomes (+0.310 vs. +0.045 µV/SD, *p* = .0395) and 416–572 ms for negative outcomes (−0.486 vs. −0.134 µV/SD, *p* = .0095). Because regression slopes carry opposite signs, steeper slopes in ADHD across both panels demonstrate amplified scaling to outcome magnitude across valences, rather than a valence- selective dissociation.

